# Horizontal transfer of a conserved *npc-2* like effector gene in rust fungi that suppresses cell death in plants

**DOI:** 10.1101/2021.08.18.456830

**Authors:** Rajdeep Jaswal, Himanshu Dubey, Kanti Kiran, Hukam Rawal, Gulshan Kumar, Sivasubramanian Rajarammohan, Rupesh Deshmukh, Humira Sonah, Pramod Prasad, Subhash C Bhardwaj, Naveen Gupta, Tilak Raj Sharma

## Abstract

ML/MD-2 is a conserved lipid/sterol-binding protein family having a role in sterol transfer and innate immunity in lower and higher eukaryotes. Here we report a genome-wide survey of this family, identifying 84 genes in 25 fungal and five oomycetes plant pathogen, having a different nutrition mode. All the fungal species were found to have varied numbers of family members, a distinctively substantial expansion of the ML gene family was observed in *Rhizophagus irregularis* (RI) with 33 genes. Our analysis also showed that NPC2 like proteins, a subfamily of ML domain superfamily, were not only restricted to animals and insect species but also present in plant fungal pathogens, including members of *Clavicipitaceae*, *Pucciniacease,* and *Tremellaceae* family. The phylogenetic analysis showed that these NPC2 like fungal proteins are more closely related to animals/insects than other fungal species. The molecular docking studies of these proteins with cholesterol and other derivatives indicate lipid-binding functional conservation across the animal and fungi kingdom. Further, the full length CDS of one of the *npc2* like genes from *Puccinia triticina* (*Pt5643*) was PCR amplified and further characterized using various studies such as qRT-PCR, expression in onion epidermal cells, *Nicotiana benthamiana* for subcellular localization studies, yeast functional complementation, and expression studies. The mRNA abundance of *Pt5643* was observed to be increased along with the infection progression and exhibits the highest expression at 5thday post-infection (dpi), suggesting its important role in the *P. triticina* infection cycle in wheat. The fluorescent confocal microscopy of transiently expressed YFP tagged *Pt5643* in onion epidermal cells and *N. benthamiana* shows its location in cytoplasm and nucleus, indicating its involvement in the manipulation of host genes. The functional complementation of *Pt5643* in *npc2* mutant yeast showed its functional similarity to the eukaryotic npc2 gene. Further, the overexpression of *Pt5643* also suppressed the BAX and H2O2 induced program cell death in *N. benthamiana* and yeast, respectively thus proving to be a novel horizontally transferred effector in rust fungal pathogens. Altogether the present study reports the novel function of fungal NPC2 like proteins playing a crucial role in host defense manipulation possibly through lipid binding/transport similar to animals.

## 1. Introduction

In nature, plants and pathogens interact with each other continuously. Both host and pathogen need myriads of small molecules like carbohydrates, proteins, lipids for various cellular processes. Out of these, lipids and sterols perform crucial roles in various processes ranging from making cell membranes to function as defense signaling molecules. Sterols constitute major components in the cell membrane of the eukaryotic cell. The presence of various sterols in different proportions constitutes the variability of cell membranes in all eukaryotes.

The effector proteins are one of the essential arsenals of fungal pathogens including rusts to disarm the host(Jaswal et al., 2020a, 2019). Most of the rust effectors are highly host-specific and do not show any similarity to known domain proteins(Jaswal et al., 2020b). Several effectors need to possess a conserved domain or fold to perform a function as an effector. The host genes’ evolutionary selection pressure causes most of the effector genes to changes their sequences. In this scenario, the pathogens change these gene sequences; however, they may maintain the minimum conserved fold to perform a corresponding biological function. Moreover, the pathogen may also generate duplicate copies to perform the function to minimize the selection pressure on the single gene. In the other case, the pathogen may also acquire certain genes from the environment to attain novel traits to deceive the host in infection pressure.

The horizontal gene transfer is also known as lateral gene transfer, is a common method of acquiring foreign genes by organisms from other species. In the case of fungal organisms, various studies have reported the presence of HGT genes donated by bacterial as well as other eukaryotic organisms like fungus, plants, and other species (Coelho et al., 2013; Fitzpatrick, 2012; Qiu et al., 2016; Slot and Rokas, 2011; Yin et al., 2016). However, relatively few studies have reported the inter-eukaryotic gene transfer.

MD-2-related lipid-recognition (ML) is one of the protein domain superfamily present in eukaryotes (plants, animals, fungi, protists) having lipid and sterol binding activity. ML superfamily includes MD-1, MD-2, NPC2 proteins, GM2 activator, phosphatidylinositol/phosphatidylglycerol transfer protein (PG/PI-TP), and mite allergen Derp2 protein, including several other proteins with unknown functions (Inohara and Nuñez, 2002). The ML Proteins generally contain a single domain (150 amino acids), N-terminal signal peptide, immunoglobulin-like antiparallel beta rich folds, and conserved cysteine residue with hydrophobic pocket present at the center suitable for binding diverse lipid/sterol molecules(Inohara and Nuñez, 2002). Crystallographic structures of various ML family proteins like NPC2, GM2-AP, and Derp2 have been studied and analyzed (Johannessen et al., 2005; Kim et al., 2007; Xu et al., 2007). The crystal structure analysis of bovine NPC2 revealed that the sterol binding site is present inside a deep hydrophobic pocket between the two beta sheets. The structure analysis also showed the binding site to be compact and small in the absence of ligand, which tends to expand after binding of a sterol ligand (Xu et al., 2007).

ML superfamily genes characterized in organisms such as insects and humans are known to perform diverse functions like providing innate immunity, chemical communication, and sterol trafficking(Frolov et al., 2003; Shi et al., 2012). The *Drosophila melanogaster* NPC2a and NPC2e proteins have a varied number of cysteine residues and bind with bacterial cell wall components *in vitro,* providing innate immunity (Shi et al., 2012). The *npc2a* and *npc2b* genes in *D. melanogaster* are similar to vertebrate *npc2* genes and regulate steroid biosynthesis and sterol homeostasis (Huang et al., 2007). In *Anopheles gambiae,* these proteins play an important role in immunity against *Plasmodium falciparum* (Dong et al., 2006). The ML protein from tobacco hornworm binds to lipopolysaccharide (LPS) and is possibly involved in LPS mediated defense signaling pathways against Gram-negative bacteria (Ao et al., 2008).

A shrimp ML domain-containing protein also binds to LPS and shows up-regulation when challenged with LPS (Liao et al., 2011). In addition to this, ML family members are also known to play an important role by acting as agonists in dengue virus infection by modifying mosquito immunity (Jupatanakul et al., 2014). One of the invertebrate ML superfamily members identified in *Ciona intestinal*is homologous to the vertebrate *npc2* gene and is induced after LPS inoculation (Vizzini et al., 2015). In the case of plant fungal pathogens, very few have been explored to characterize the functions of this superfamily. A*npc1* like gene has been reported to perform sterol trafficking in the *Fusarium graminearum* (Breakspear et al., 2011). *Metarhizium robertsii* an insect fungal pathogen, has acquired *npc2-* like gene from its insect host in the course of evolution through horizontal gene transfer (Zhao et al., 2014). No other study has reported the function of this superfamily in the plant-fungal pathogens In the present study to explore the status of the ML superfamily, the genome-wide identification method has been employed in 25 fungal and five oomycetes species having different modes of nutrition. We have compared these proteins at the sequence level, analyzed the chromosome location, and identified the phylogenetic relationship across eukaryotes (fungi, plants, animals, and protists). The comparison of structures of proteins at the tertiary level three and molecular docking was done to find out the possible interacting partners. Selection pressure studies were also performed. Expression patterns were also studied using the SRA NCBI database as well as the real-time PCR. Subcellular localization was also performed to predict the location of one of the *P. triticina npc2-* like genes.

The functional similarity of the rust *ml* gene *Pt5643* was also checked using complementation assay in the *npc-2* mutant of the *Saccharomyces cerevisiae*. Moreover, *in-planta and S.cereviseae* expression of *Pt5643* was done to evaluate its capacity as an effector. This study provides the first report of the presence of *npc2* like genes in rust fungi and further its role as an effector. Moreover, the study also provides a baseline for exploring the ML superfamily in eukaryotes Additionally, it can also be utilized to elucidate the evolutionary relationship of the ML gene family across plant fungal pathogens and predict its functional importance as an effector.

## 2. Materials and methods

### 2.1. Identification of ml/md-2genes across fungal pathogens

The fungal proteomes were downloaded from the UniProt database (http://www.uniprot.org/help/uniprotkb) and NCBI (https://www.ncbi.nlm.nih.gov). The predicted protein sequences of *Puccinia triticina* and *Puccinia striiformis* used for this study were taken from the data generated by Kiran et al., 2016 and 2017 (Kiran et al., 2017, 2016). All the putative ML/MD-2 domain-containing proteins were retrieved from the Interpro database (https://www.ebi.ac.uk/interpro/entry/IPR0031672). The psi-blast program was used to perform genome-wide identification using downloaded ML/MD-2 proteins (IPR033916) as a query against all the organisms (Altschul et al., 1990). Target proteins with BLAST hit (E-value<-5) were selected and subjected to NCBI-CD search (https://www.ncbi.nlm.nih.gov/Structure/cdd/wrpsb.cgi) and InterPro database (http://www.ebi.ac.uk/interpro/search/sequence-search) for conserved domain analysis. Redundant and nonspecific hit-containing proteins were filtered out, and proteins with ML/MD-2 domain-containing proteins were selected as candidate proteins.

### 2.2. Secretory protein prediction, Subcellular localization, and physicochemical properties analyses

Secretory protein analysis was done by using SignalP 3.0 for signal peptide analysis (http://www.cbs.dtu.dk/services/SignalP/). TargetP1.1 and CELLO were used to analyze the subcellular localization of genes. (http://www.cbs.dtu.dk/services/TargetP/), (http://cello.life.nctu.edu.tw/). The TMHMMv2.0 was used for finding transmembrane-containing proteins. (http://www.cbs.dtu.dk/services/TMHMM/). GPI Modification Site Prediction in Fungi was predicted using big-PI Fungal Predictor (http://mendel.imp.ac.at/gpi/fungi_server.html). The amino-acid and atomic compositions, theoretical pI, the composition of amino acids, the molecular weight of all ML/MD-2 proteins was calculated using the ExPASy server’s ProtParam and Compute pI/Mw tool. (http://web.expasy.org/protparam/), (http://web.expasy.org/compute_pi/). All the tools were run using default parameters.

### 2.3. Chromosomal localization and gene structure analysis, Motif and Domain analysis

The chromosomal location of the *ml* genes was determined for fungal pathogens having chromosomal data in NCBI using Mapchart 2.2. BLASTn was done to retrieve the specific location of genes for the corresponding organism. Gene structures of *ml* genes were generated using GSDS (http://gsds.cbi.pku.edu.cn/); motifs were predicted using the MEME program (http://meme.nbcr.net/meme/cgi-bin/meme.cgi). The Parameter selected for the analysis was as follows: total motif predictions were restricted to 10, and the size of the motif was given from 4-50 amino acids. The full-length ML/MD-2 proteins were annotated first using CD-search for different domain identification. The representing structures of various domains were designed using IBS 1.0 (http://ibs.biocuckoo.org/).

### 2.4. Protein Structure prediction and docking analysis

3D Structures of all the secretory proteins were predicted using I-Tasser software and further refined by modrefinner(Xu and Zhang, 2011; Yang et al., 2015). Ramachandran plot analysis was done using Rampage software, showing > 80% amino acids in the core region for all the structures (http://mordred.bioc.cam.ac.uk/~rapper/rampage.php). Final validation for all the structures was done using tools available on the SAVES server (https://services.mbi.ucla.edu/SAVES/). Docking analysis was done using the Docking Server (Bikadi and Hazai, 2009). Ligands were retrieved from PubChem in sdf format and changed in pdb format with the help of an open babel tool. All the structures were converted into energy minimized states using Hamiltonian-Force Field-MMFF94x (Halgren, 1996). Gasteiger partial charges were added to ligand atoms. AutoDock tool was used to add Kollman united atom type charges, essential hydrogen atoms solvation parameters.

### 2.5. Multiple sequence alignment and phylogenetic analysis

The sequence alignment of ML/MD-2 proteins was done using MAFFT software (online version) (http://www.ebi.ac.uk/Tools/msa/mafft/). For comparative analysis, proteins from major host plants of different pathogens and few model species (Rice, Wheat, Tomato, Potato, Maize, Soybean, and Human, insects, protists) were included in the study and aligned. For sequence alignment, the ML/MD-2 domain region was extracted using python script for all the proteins (including host plants and model species). For phylogenetic analysis, full-length proteins were aligned using MAFFT, model testing was done using MEGA7.0 (Kumar et al., 2016), and the best-fitted model was considered for the tree construction. The phylogenic tree was constructed in MEGA7.0 with the maximum likelihood (ML) method with 1000 bootstrap replicates. A total of 68 different eukaryotes (Fungi-22, Plants-6, oomycetes-5, protists-28, animals-5) were used in the analysis to infer the phylogenetic relationship of ML proteins among these species.

### 2.6. Selection pressure analysis of the *ML* gene family

The nonsynonymous substitutions (Ka) and synonymous substitutions (Ks) were utilized to analyze section pressure of the *ML* genes (Li et al., 1981). DNASP5.0 software was used to analyze the Ka/Ks ratios between the genes (Librado and Rozas, 2009). Nucleotide sequences (CDS) were aligned first and used as a query. Results were analyzed on the presumption that purifying selection causes values less than 1 (Ka/Ks < 1), and positive selection leads to a higher value than 1(Ka/Ks > 1) (Juretic et al., 2005).

### 2.7. Plant materials, pathogen inoculation, tissue harvest, RNA extraction and real-time quantitative PCR

Susceptible wheat variety (Agra Local) was used as a host for infection of *P. triticina*, *P. striiformis,* and *B. graminis*. Uninfected (Control) and infected leaves were harvested at various time intervals (0,3,5,7 9 and 14 dpi), respectively, immediately frozen in liquid N2 and stored at −80°C for further analysis. Total RNA was isolated from mirVana™ miRNA Isolation Kit, Thermo Fisher Scientific (USA).

First-strand cDNA synthesis was done from RNA using the iScript™ (Biorad) USA reverse transcription kit with approximately 1 μg RNA according to the manufacturer’s instruction. For qRT-PCR, primers were designed using Oligocalc software and with amplicon lengths of 120–250 bp. The qRT-PCR was done on 7500 ABI Real-time PCR systems (Applied Biosystems, United States) using Taq™ Universal SYBR® Green Supermix(Biorad) USA. The relative expression was calculated by the 2^−ΔΔCt^ method(Livak and Schmittgen, 2001). The *PtActin* and *PtGAPDH* were used as internal controls.

### 2.8. Cloning, In-vitro, In-planta, and yeast expression

Amplification of the coding sequence (CDS) *Pt5643* gene (*P. triticina npc2* like gene) was done using full length gene-specific primers with BamHI (forward primer) and NotI (reverse primer) restriction sites and cloned into the pGEM-T easy vector. Confirmation of the clone was done through PCR, restriction digestion, and sequencing. The confirmed fragment was further mobilized into the pET 29a (+) vector and transformed into BL21 *E.coli* expression strain; induction for invitro expression was done by using 0.1 mM IPTG at 37 °C and analyzed on the 12% SDS-PAGE. For *In-planta* expression and subcellular localization experiments, full length CDS (without signal peptide coding region and stop codon) of *Pt5643* and BAX gene (full length CDS PCR amplified from mouse cDNA) was cloned using gateway cloning vector pENTER and further mobilized into the destination pGWB408 for overexpression. For subcellular localization, the pENTER_*Pt5643* clone was used for mobilizing the *Pt5643* in the pGWB441. Dr. Tsuyoshi Nakagawa’s lab generously provided the binary plant transformation vector pGWB408 and pGWB441. For expression in the yeast, the pENTER_*Pt5643* clone was used for mobilizing the *Pt5643* in the pYESDEST52 gateway destination vector.

### 2.9. Subcellular localization of Pt5643 gene

For subcellular localization, onion epidermal cells, particle bombardment system, and a *Nicotiana benthamiana* agrobacterium mediated transient expression methods were used. The confirmed pGWB4441_*Pt5643* construct (YFP) with the C-terminal fluorescent tag and an empty vector containing only fluorescent tag was bombarded separately on onion peels containing MS plates using the PDS1000/He system at 1100 psi (BioRad, USA). After 36 hrs, the samples were visualized under Zeiss LSM 880 with laser scanning confocal microscope (Carl Zeiss Microscopy, USA).

To confirm the subcellular localization of *Pt_5643*, 0.5 optical density (OD) agrobacterium (strain GV3101) culture containing *Pt_5643*_pGWB441 and empty vector was infiltrated in separate *N. benthamiana* leaves and analyzed after three days at the infiltrated areas at 514 nm in the confocal microscope.

### 2.10. Functional complementation assay in npc2 yeast mutants

For function complementation assay, the wild-type yeasts BY4741 and the *npc2* mutant (YDL046W) were used. The npc2 mutant competent cells were transformed with pYESDEST52 and pYESDEST52_*Pt5643* construct separately and further selected on the uracil dropout medium. The functional complementation was analyzed on 100mM acetic acid plates by spotting assay (1-1/1000 dilutions) as the npc2 mutants are resistant to acetic acid, whereas the wild type is susceptible. The phenotypes were analyzed after 3-4 days.

### 2.11 Bax and H2O2 induced cell death in transient expression in *N. benthamiana* and *S. cerevisiae*

The pGWB408 empty vector, pGWB408_BAX, and pGWB408_*Pt5643* constructs were transformed in *Agrobacterium tumefaciens* GV3101 competent cells and further confirmed by PCR. *N. benthamiana* plants grown up to 3-4 weeks were used agroinfiltration. The cultures were adjusted to an optical density (OD) of 0.5 and suspended in MES buffer containing 10 mM MgCl2, 10 mM 2-(N-morpholino) ethanesulfonic acid (MES), pH 5.6, and 100 μM acetosyringone. The BAX construct was infiltrated 0, 12, and 24 hrs post *Pt5643* construct infiltration in different spots. The phenotype was observed after 5-8 days of infiltration. The H_2_O_2_ induced oxidative stress hijacking phenotype analysis was done using 5mM H_2_O_2_ containing YPD plates. Approximately 1/1000 cells were plated from the OD 1.O of yeast cells for *npc2* mutant cells, wild type (BY4741), and pYESDEST52_*Pt5643* containing *npc2* mutants grown in YPD broth medium. The results were analyzed after 3-4 days.

## 3. Results

### 3.1 Genome-wide Identification of ML domain superfamily and physiochemical properties analysis

To identify ML/MD-2 proteins from various fungal species, interpro-id IPR0031672 specific for ML/MD-2 proteins was used as a query against the proteome of the target organisms (Supplementary file S1). A varied number of proteins from different organisms were found (Figure 1A). Most of the species had one to three copies of *ml* gene except *B. graminis* with five and *C. purpurea* with four *ml* gene copies each. Apart from these two pathogens, the highest numbers of genes (33) were found in *R. irregularis*. Further, these genes were subjected to NCBI CD-search to verify their coding region for the ML domain (Supplementary file S2). The deduced sequences varied between 139 amino acid residues (SCD8PQS8) to a maximum of 1071 amino acid residues (TCL8WPE2). Cysteine content analysis for ML protein showed the highest cysteine content in *the Puccinia triticna* Pt5643 gene, while CHM2TBX0 had the lowest cysteine content (Supplementary file S3, S4, and S5).

**Fig. 1.**
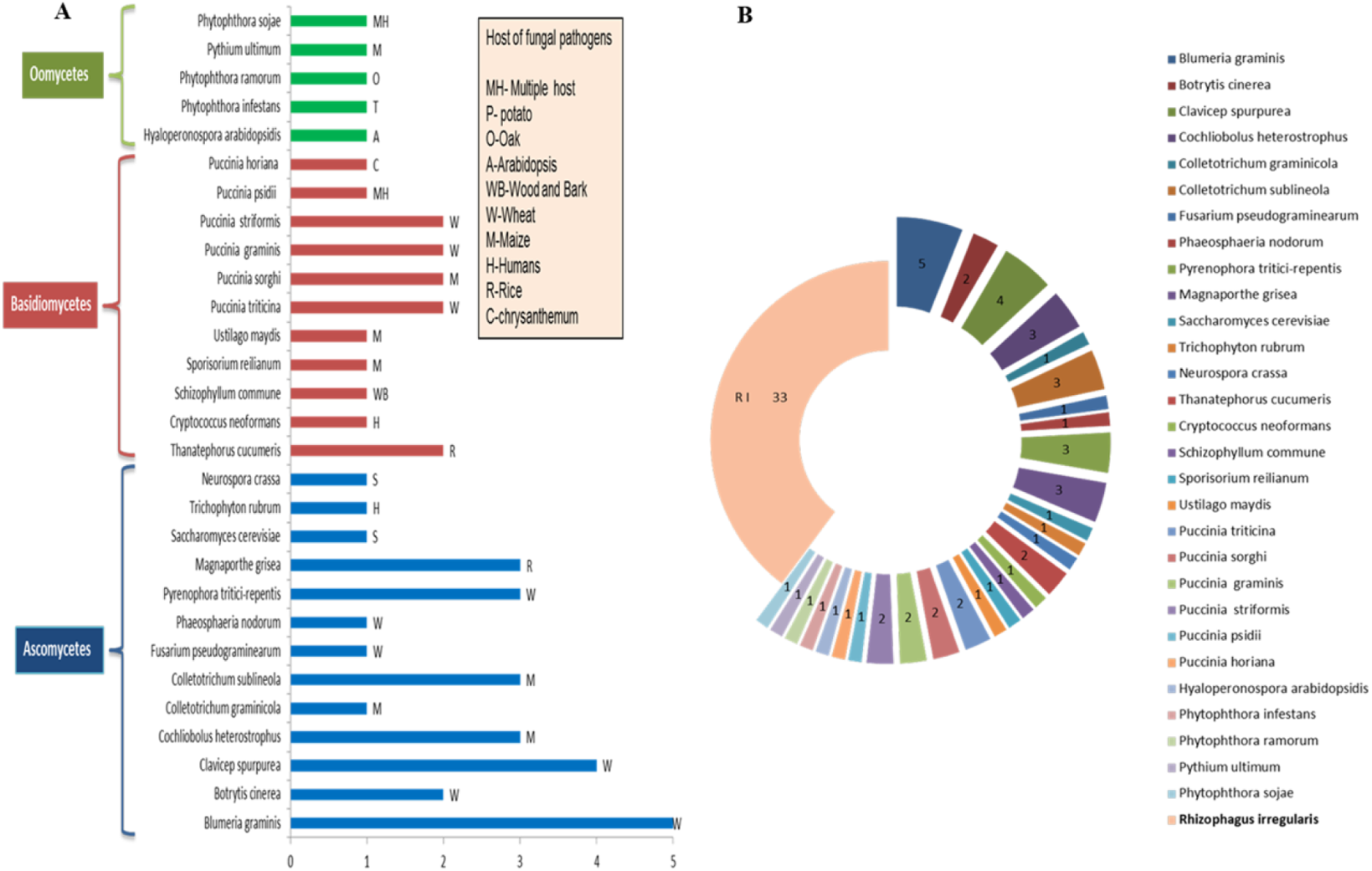
Genome-wide identification of *ml* gene family in various fungal species. **(A)** ML domain proteins identified in major fungal and oomycetes species. Among biotrophs, *Blumeria garminis* and *Claviceps purpurea* had the highest ML proteins 5 and 4, respectively, while other species (hemibiotrophs, necrotrophs, saprotrophs and oomycetes mostly enode1-3 copies. **(B)** Comparison of identified ML proteins of various fungal species with mycorrhizal fungus *R. irregularis* (RI)*., the* highest number of ML domain proteins (33) were found in RI, suggesting an expansion of protein domain family.

### 3.2 Protein domain, exon/intron structure, and conserved motifs analysis

The NCBI CD search and Pfam were used to identify the conserved domain present in ML proteins. The results revealed that ML superfamily proteins are enriched with a single ML domain, while some possessed other multiple domains apart from the ML domain (Figure 3). Proteins containing a single ML domain (number) were most abundant, with a minimum length of 46 amino acids for *B. graminis* (BGN1JH01) and a maximum length of 138 amino acids for *P.horiana* (PH14254). The second most abundant group was proteins containing ML domain at N-terminal and TRP-like a dominant domain at C-terminal ends. Proteins exclusively present in oomycetes comprised the third group with two additional domains, the Cathepsin propeptide inhibitor domain at N-terminal and Peptidase C1A papain along with the small-sized ML domain at the extreme C terminus. The fourth group consisted of two proteins (TCL8WM75, TCL8WPE2) from *Thanatephorus cucumer*, having three different domains, the SKBP1/BTB/POZ domain and the Nucleophile aminohydrolases and Carbohydrate-Binding Module family (CBM) domain respectively. The exon/intron structural arrangements and their number were similar for species from the same class (Data not shown) Interestingly most of the ascomycetes species had three exons and two introns, except *Colletotrichum sublineola* (CSA0A066WWY1) and *Saccharomyces cerevisiae* (SC02278) which were intronless. The motif size identified by MEME was 4 to 50 amino acids long. Similarity within the motifs across fungal and model host ML proteins showed three major motifs for these proteins (Figure S4). The first group represented the NPC2 domain present in basidiomycetes, while the second motif was represented by PG/PI group proteins, causing the majority of single domain ML (PG/PI) proteins to fall in this group. The third group comprised the same NPC2 domain; however, both the ascomycetes, as well as basidiomycetes proteins, were present in this group.

**Fig. 3.**
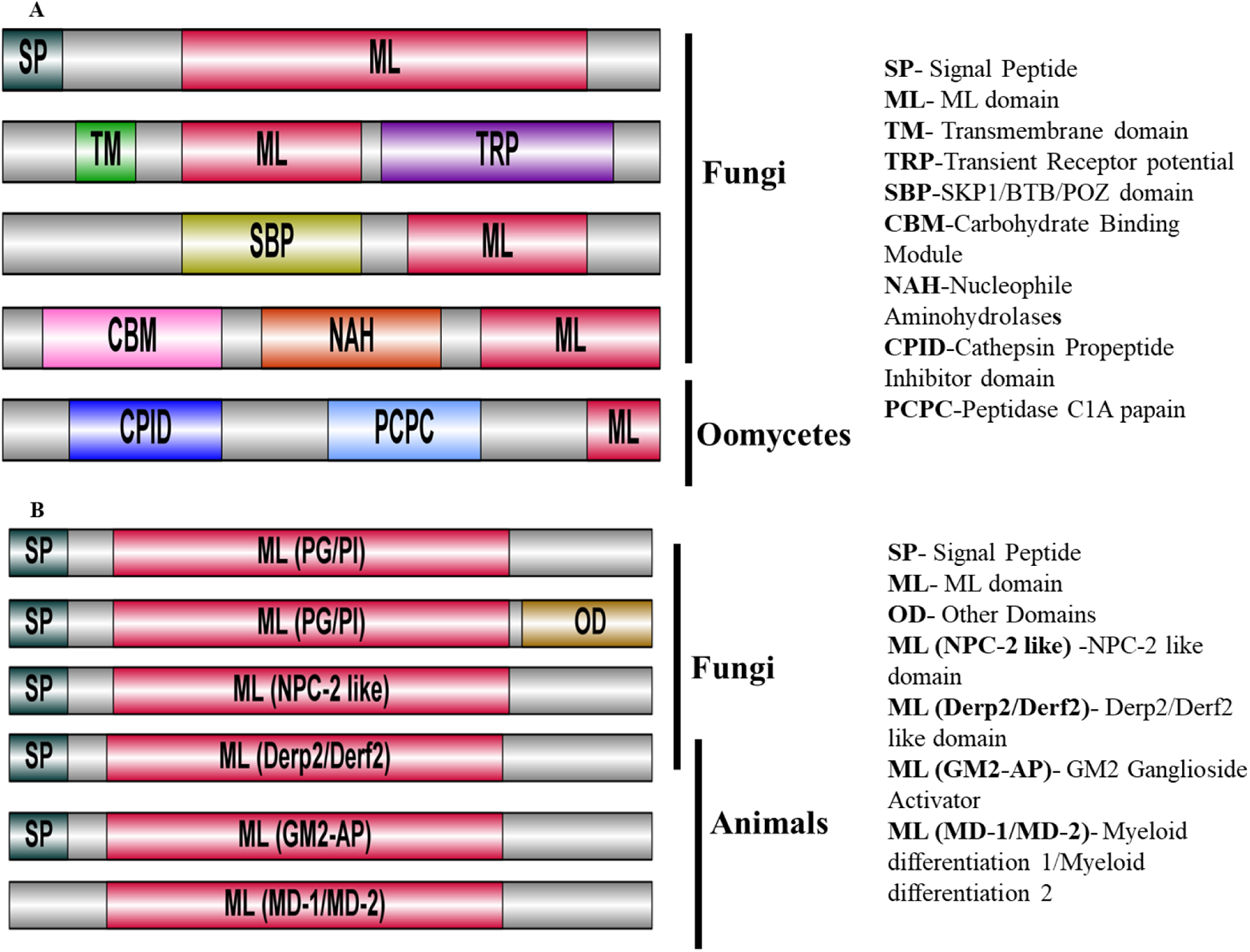
Domain organization of ML protein family. **(A)** Domain organization of ML and associated domain in fungal and oomycetes proteins: the size of the domains used only for representation and does not represent the actual size. Most fungal species contain a single ML domain; however, few members from ascomycetes have double domain annotated as TRP (Transient Receptor Potential), while oomycetes ML members contain Cathepsin Propeptide Inhibitor domain (CPID) and PCPC-Peptidase C1A papain (PCPC). **(B)** Domain organization of ML proteins for host and model species: most of the hosts (animal and plant ML proteins) have a single ML domain, unlike fungal and oomycetes ML proteins.

### 3.3 Sequence alignment and characterization of ML superfamily proteins

Multiple sequence alignment of the identified ML proteins showed very high diversity among and across the fungal species was observed, except in the domain region that was relatively conserved (Figure S1A). All ML proteins generally had four conserved cysteine and basic residues (Lysine and Arginine) in the domain region, which are essentially required for disulfide linkage and making a hydrophobic pocket for ligand binding(Gruber et al., 2004). Further, the alignment of conserved domain regions revealed the clustering of all of the sequences into four major groups (Figure S1B). Group one included proteins of the PG/PI category present in plants and fungal species only. Group two included proteins from humans, mice, and other animals annotated as GM2-AP of the ML family. The third group consisted of NPC-2 like proteins from drosophila, humans, *Clavicipitaceae*, *Tremellaceae,* and rust fungal species. The fourth category was formed by MD-1/MD-2 proteins present only in animals. Except for a few species, most of them were grouped as expected. The identified ML proteins of *Clavicipitaceae* and *Tremellaceae* clustered along with the NPC2 and PG/PI proteins, which are from the third and first groups, respectively. However, *Puccinia sp* ML proteins specifically clustered only with group three NPC2 proteins. These results were quite interesting to observe the unusual presence of NPC2 like domain within the fungal species.

So to investigate the origin of *npc2-*like genes in *P.triticina* and other species with NPC2 domain, BLASTp analysis was performed using the NCBI non-redundant (nr) database (1.0e-5, and percentage identity of 30%) against Pt5643, a *P. triticina* NPC2 like protein. As expected, top hits for the *Pucciniaceae* family were achieved, followed by hits with species *Kockovaella imperatae*, *Naematelia encephala,* and multiple members of *Kwoniella,* all from family fungi/*Tremellaceae*, few hits with *Culex quinquefasciatus* and *Tetranychus urticae* from family insect/dipteral were also observed. (Supplementary file S7). The results were reconfirmed by performing tBLASTn analysis using *Pt5643* (Supplementary file S8). Additionally, we also used *Kockovaella* sp and Culex ML protein as a BLAST query against the NCBI nr database as well as exclusively against fungal taxa. We found *Puccinia*ceace and *Tremellaceae* family members as top hits (Supplementary file S9).

Further, BLASTp using *Pt5643* as a query against the Insect Base database was performed. Major hits with high e-value found were from *C. quinquefasciatus, Anopheles,* and *Mayetiola destructor* all belonging to *Diptera* (Supplementary file S10 & S11). Apart from *Pt5643* as query, BLAST analysis using previously reported horizontally transferred from *Metarhizium* and *Claviceps npc2-* like genes (as a query) with similar parameters demonstrated that NPC2 like proteins are well present in other species like *Pochonia chlamydosporia* and *Ustilaginoidea virens* from class *Clavicipitaceae* (Supplementary file S12). Additionally, Pfam, Interpro, and SMART databases were also searched for ml proteins and annotated with CD-search to explore the possibility of finding any novel NPC2 like proteins; we did not find any other fungal NPC2 like proteins in these databases except for the above-reported species.

### 3.4 Phylogenetic studies and selection pressure analysis

Phylogenetic analysis was performed using the IQ-TREE server (Trifinopoulos et al., 2016). The ML proteins of fungal and oomycetes species identified in this study along with previously known fungal ML proteins from family *Clavicipitaceae*, as well as from various other eukaryotes (plants, animals, insects, protists (retrieved from Interpro) were combined to compare and infer the existing evolutionary inference. The analysis showed oomycetes ML proteins as a separate clade pointing towards its unrelatedness from fungal species (Fig 2). According to the previous study (Inohara and Nuñez, 2002), fungal ML proteins contain PG/PI domain only. The results showed PG/PI domain proteins cluster in two separate groups, one containing single PG/PI domain-containing proteins while other TRP domain proteins associated with PG/PI domain. The latter group was found specifically present in ascomycetes ML proteins. In the case of animals, three major subclusters were formed belonging to the class, GM2-AP, MD-1-MD-2, and NPC2, respectively.

**Fig. 2.**
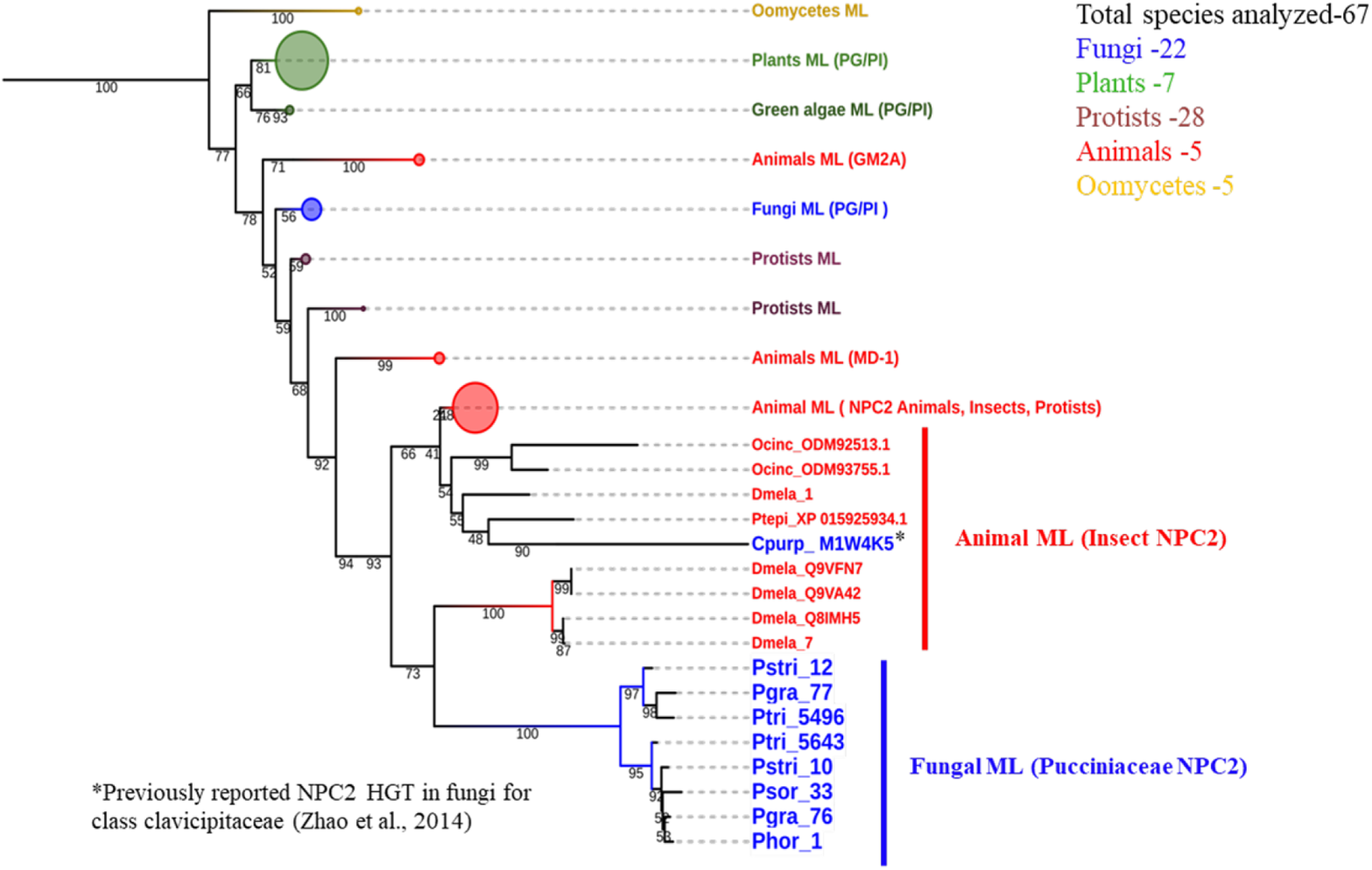
Phylogenetic distribution of ML domain protein family. Phylogenetic relationship of ML proteins in plants, animals, insects, protists and fungi: the proteins clustered in four subclasses: ML Animals (MD-1,GM2A, NPC2), plant, green algae and fungi (PG/PI), Fungi HGT events (NPC2 (*Pucciniaceae* and *Clavicipitaceae*). For tree construction, maximum likelihood method with 1000 bootstrap runs was used). The tree was generated using IQ-TREE software.

The ML proteins from *Clavicipitaceae* (*C. purpurea* and *Metarhizium robertsii*) have already been reported to cluster with animal NPC2 like proteins (Zhao et al., 2014). In the context of this, the results additionally showed ML proteins from other multiple *Metarhizium* spp to cluster with animals NPC2 proteins. Surprisingly we also found that NPC2 like proteins of other members like *P. chlamydosporia* and *U.virens* also clustered with animals NPC2 proteins (Fig 2). Apart from *Clavicipitaceae* (ascomycetes) members, various rust fungal species (basidiomycetes) also clustered with animals NPC2 like proteins, however, forming separate subgroups from both animals NPC2 and *Clavicipitaceae* suggesting its uniqueness from these species. Though, fungal NPC2 proteins seemed to show more similarity with animals and insects than the rest of the fungal ML proteins (Fig 2). In contrast to so much diversity within the fungal ML proteins, plant PG/PI proteins were present only in one separate cluster supporting the previous reports (Inohara and Nuñez, 2002). Separately, the Ka/Ks ratio analysis for orthologous ML genes was also performed. Results demonstrated that most of the ML genes are under purifying selection except two genes from *B. graminis*, one from *P. striformis,* and *R. irregularis* possessed Ka/Ks >1 and therefore might be under possible positive selection (Table 1).

**Table 1.** Positively Selected gene pairs of ML gene family in fungal species.

| **Gene1** | **Gene2** | **Ks** | **Ka** | **Ka/Ks** |
| --- | --- | --- | --- | --- |
| *BGNIJH01* | *BGN1JF5* | 1.11 | 2.38 | 2.13 |
| *BGAoA078N* | *BGN1JF5* | 2.47 | 3.10 | 1.26 |
| *PSTRI10* | *PSTRI12* | 1.29 | 1.72 | 1.33 |
| *U9SUS7_RI15* | *U9SSM5_RI4* | 0.0656 | 0.1035 | 1.57 |

### 3.5 Secretory proteins analysis, subcellular localization, and genomic distribution of ml genes

To identify signal peptide and putative subcellular location of identified ML proteins, CBS prediction Servers software was used, and results were summarized in Supplementary Table 1. EffectorP marked 32 proteins out of 84 as putative effector molecules. Overall, these *insilico* analyses showed that ML proteins are diverse in location and can be found in extracellular secreted form or membrane-bound form to different organelles. The chromosomal location of ML family genes was analyzed by using mapchart software (Figure S5).

### 3.6 3 Dimensional structure prediction and molecular docking studies

To analyze the interaction of *ML* proteins with the various lipid and sterols, tertiary structures of all ML proteins were examined using various bioinformatics tools. Proteins that were extracellular secretory in prediction were used further for the 3D structure predictions (Figure 4 and Figure S6). For Molecular docking, various sterols and lipids were used, but cholesterol and lanosterol were used for the initial screening of ML proteins. Out of the analyzed species, *B. graminis*, *C. purpurea*, *P. triticina, P. sorghi, P. striiformis, and R. irregularis* showed positive docking using cholesterol, lanosterol, and various ligands (Table 2).

**Fig. 4.**
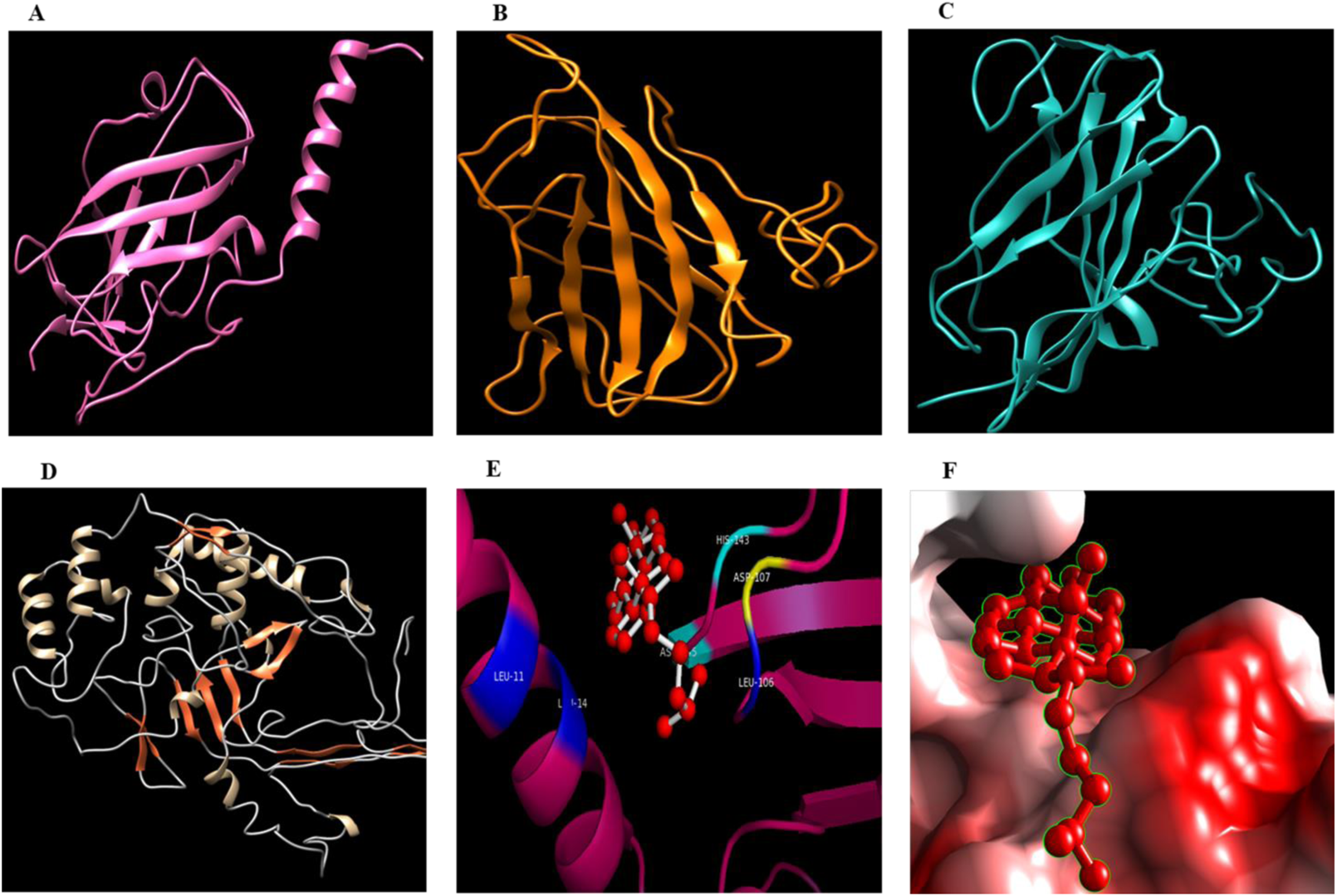
The 3D Structure prediction and molecular docking analysis of ML domain proteins. Few structures were used here for representing the structure of ML proteins. (A) NPC2 like protein of *P.triticina*. Figure (B) PG/PI protein of *M.oryzae*. (C) PG/PI TRP protein of *S.reilianum.* D) ML protein of *P.ramorum*. (E) & (F) Molecular Docking of Pt5643 (NPC2 like proteins) of *P.triticina* with cholesterol. The cholesterol showed the highest docking score with Pt_5643. Ribbon and surface forms of 3D structures were used to highlight the interactions. 3D structure of all proteins is available in Figure S5 (Supplementary data).

**Table 2.** Molecular docking analysis of selected ML domain proteins from fungal species with various ligands.

| **Organism Name** | **Gene Name** | **Ligand** | **Est. Free Energy of Binding kcal/mol** | **Total Intermolecular Energy(kcal/mol)** |
| --- | --- | --- | --- | --- |
| *P. triticina* | *Pt_5643* | Zymosterol | -6.43 | -7.60 |
| *P. triticina* | *Pt_5643* | Cholesterol | -6.52 | -7.79 |
| *P. triticina* | *Pt_5643* | GP | -4.27 | -6.20 |
| *P. triticina* | *Pt_5643* | Lanosterol | -6.43 | -7.93 |
| *P. triticina* | *Pt_5643* | Lathosterol | -6.30 | -8.10 |
| *P. triticina* | *Pt_5643* | IPP | -2.14 | -7.81 |
| *P. triticina* | *Pt_5643* | Fungisterol | -6.49 | -8.05 |
| *P. triticina* | *Pt_5643* | Sequalene | -4.98 | -8.84 |
| *P. triticina* | *Pt_5643* | FPP | -4.25 | -7.02 |
| *P. triticina* | *Pt_5643* | Stigmast -7 enol | -0.27 | -2.87 |
| *P. triticina* | *Pt_5643* | P3P | -1.74 | -4.07 |
| *C. pupurea* | *CpM1WA48* | Ergosterol | -10.03 | -11.15 |
| *C. pupurea* | *CpM1WA48* | Sequalene | -8.35 | -11.85 |
| *C. pupurea* | *CpM1WA48* | PS | -4.22 | -6.94 |
| *C. pupurea* | *CpM1WA48* | FPP | -7.35 | -9.89 |
| *C. pupurea* | *CpM1WA48* | P3P | -5.51 | -8.52 |
| *C. pupurea* | *CpM1WA48* | Cholestrol | -9.79 | -11.26 |
| *C. pupurea* | *CpM1WA48* | 24 ethldiene lophenol | -10.57 | -11.89 |
| *C. pupurea* | *CpM1WA48* | Lanosterol | -10.12 | -11.45 |
| *U. maydis* | *UMo4733* | Desmosterol | -7.24 | -8.56 |
| *U. maydis* | *UMo4733* | Fecosterol | -6.67 | -8.30 |
| *U. maydis* | *UMo4733* | Ergosterol | -6.43 | -7.86 |
| *U. maydis* | *UMo4733* | Lanosterol | -5.43 | -8.41 |
| *U. maydis* | *UMo4733* | P3P | -2.68 | -4.82 |
| *P. sorghi* | *Psorghi_33* | Cholesterol | -7.44 | -8.96 |
| *P. sorghi* | *Psorghi_33* | Lanosterol | -5.95 | -10.49 |
| *P. sorghi* | *P sorghi_33* | Fungisterol | -8.05 | -10.61 |
| *P. sorghi* | *Psorghi_33* | Ergosterol | -6.45 | -7.88 |
| *P. sorghi* | *Psorghi_33* | FPP | -2.54 | -5.79 |
| *B. graminis* | *BGAoAo78N1M3* | Cholesterol | -6.49 | -9.94 |
| *B. graminis* | *BGAoAo78N1M3* | Lanosterol | -5.95 | -10.49 |
| *B.graminis* | *BGAoAo78N1M3* | Fungisterol | -8.05 | -10.61 |
| *R.irregularis* | *U9Uo13_RI* | Cholesterol | -9.05 | -10.68 |
| *R.irregularis* | *U9Uo13_RI* | Brassicasterol | -9.04 | -10.56 |
| *R.irregularis* | *U9Uo13_RI* | Fungisterol | -9.46 | -11.11 |

\*GP- Geranyl pyrophosphate, IPP- Isopentenyl pyrophosphate, FPP-Farnesyl pyrophosphate

Since the ML protein superfamily has been reported to bind with sterols and lipids in various studies (Xu et al., 2007; Zhao et al., 2014), therefore, we focused the study on major sterols like cholesterols, ergosterol, and their homologs, plant-related sterols, Phosphatidylinositol/Phosphatidylglycerol(PI/PG), including other molecules primarily found in cholesterol biosynthesis and various other major sterol synthesis pathways of fungi. Amongst all the pathogens used in the study, *R. irregularis, B. graminis,* and *C. purpurea* showed the highest number (33, 5, and 4 respectively) of ML genes. One gene, each form *B. graminis* and *C. purpurea* showed positive docking out of all the genes used for docking with various sterol ligands. Further, randomly picked 5 ML genes were used for structure prediction and docking analysis for *R.irregularis;* All of these genes showed positive interaction with major sterols. One out of the two *P. triticina* ML proteins (the only member from rust fungi (*Pt5643*) used initially in this study) showed positive docking results (Table 3).

Further to gain confidence with the results found in *P. triticina*, other *Puccinia* species viz. *P. graminis, P. striformis, P. sorghi*, and *P. horiana* were included in the study. Docking analysis revealed a total of three genes from *Puccinia* species, one each from *P. triticna, P. striiformis,* and *P. sorghi* showing positive interaction (Table 3). None of the *ml* genes from *P. graminis* and *P. horiana* showed any interaction with the sterol ligands. It was interesting to note that all the pathogens showing positive docking interactions were biotrophic in nature. Further, analysis of the amino acid residues involved in interacting with sterols revealed hydrophobic residues to be among the top interacting amino acids followed by polar and charged residues. (Supplementary file S5)

### 3.7 Transcript abundance analysis and orthologous gene identification

The presence and abundance of ML genes in different tissue and infection stages at different time intervals were analyzed using NCBI SRA BLAST search. CDS of selected genes were subjected to SRA BLAST and significant hits (number) with E-value−5, bit score 100 were converted into reads per million (RPM)(https://www.ncbi.nlm.nih.gov/news/11-19-2013-SRA-BLAST). Converted RPM values were plotted as a heatmap using the heat mapper software (Figure 8a). The highest hits were represented by *P. striformis* genes followed by genes *from B. graminis*. The reads abundance of *P.stri*12 (425.8 RPM) and BGA0A078N1M3 (805.90 RPM) were highest in the latter stages of infection and showed positive docking interactions. Initial infection stages (6h,12h, 24h) depicted low reads in BGA0A078N1M3, while a substantial increase was observed at 48h and being highest at 5 dpi.

**Fig. 8.**
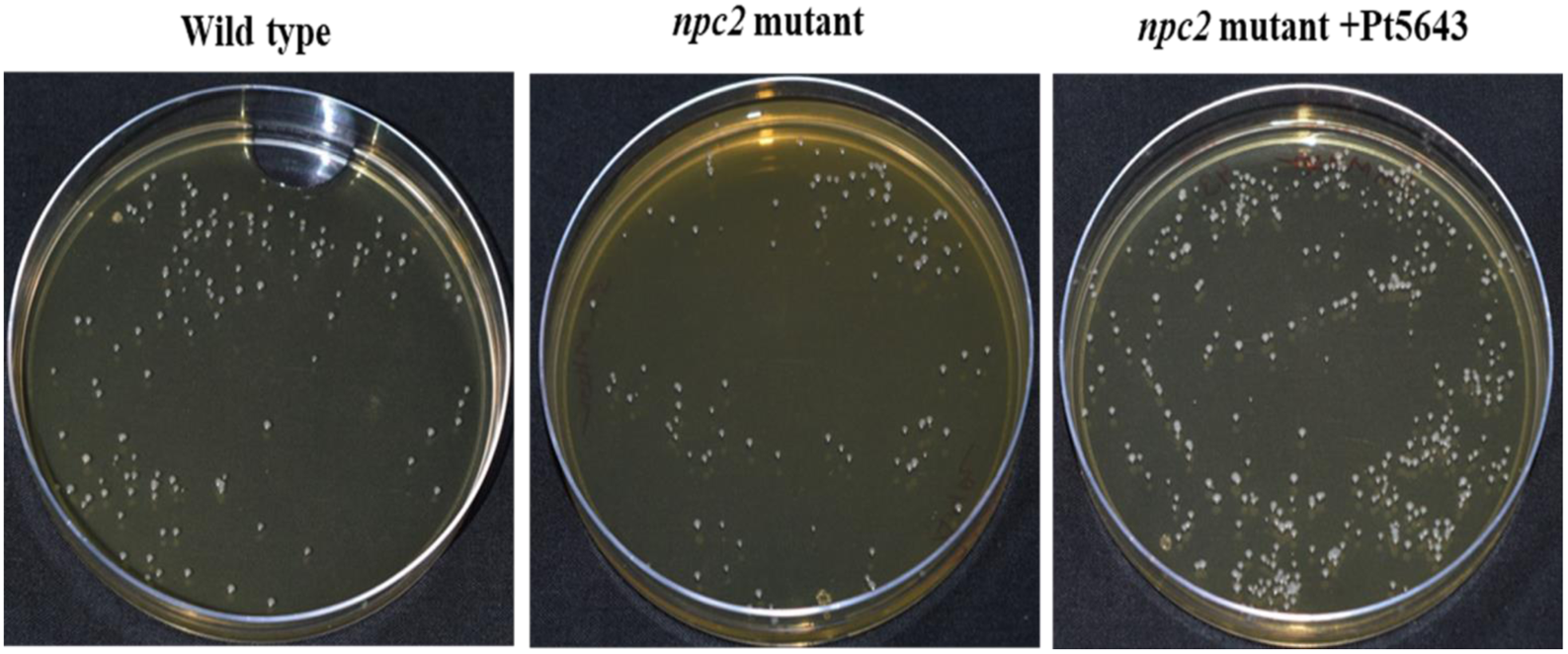
H_2_O_2_ induced cell death suppression assay of *Pt5643* gene in *S. cerevisiae*. Cells with 0.0001 each for wild type, npc2 mutant and *Pt5643* containing npc2 cells were plated on 5mM H2O2 containing YPD plates. Higher numbers of colonies approximately (324)were rescued for *Pt5643* containing npc2 cells than wild type (123) and npc2 mutant alone (90).

Orthologous gene identification across the selected pathogens (*B. graminis*, *C. pupurea*, *P. triticina*, *M. oryzae*, *R.irregularis*, *C. sublineolum*) showed very few sequences to have orthologue due to high diversity among the species. However, the orthologous gene identification within *R.irregularis* showed numerous genes to be paralogous, probably as a result of duplication events (Supplementary File S13)

### 3.8 Cloning, In vitro expression, qRT-PCR analysis, and subcellular localization

The full length ORF of the gene (*Pt5643*) with an expected size of 570 bp was amplified using gene-specific primers (Figure 5A) and clone in the p-GEMT vector to confirm the identity of the domain (NPC2 like domain) and overall sequence of the *Pt5643* gene. The sequence of *Pt5643* gene found after sequencing was again subjected to BLAST in a similar procedure previously followed for finding the similarity among other *npc2* genes of other species. Similar results were found in the case of predicted gene *Pt5643* of the *P. triticina* genome. The SDS-PAGE of the expressed protein showed an expected size of approximately 32kDa (Figure 5B). The qRT-PCR analysis was done for the gene showing positive docking interaction; however, only *Pt5643* showed a single band using RT-PCR primers and was used as a candidate for expression analysis (Figure 5C)

**Fig. 5.**
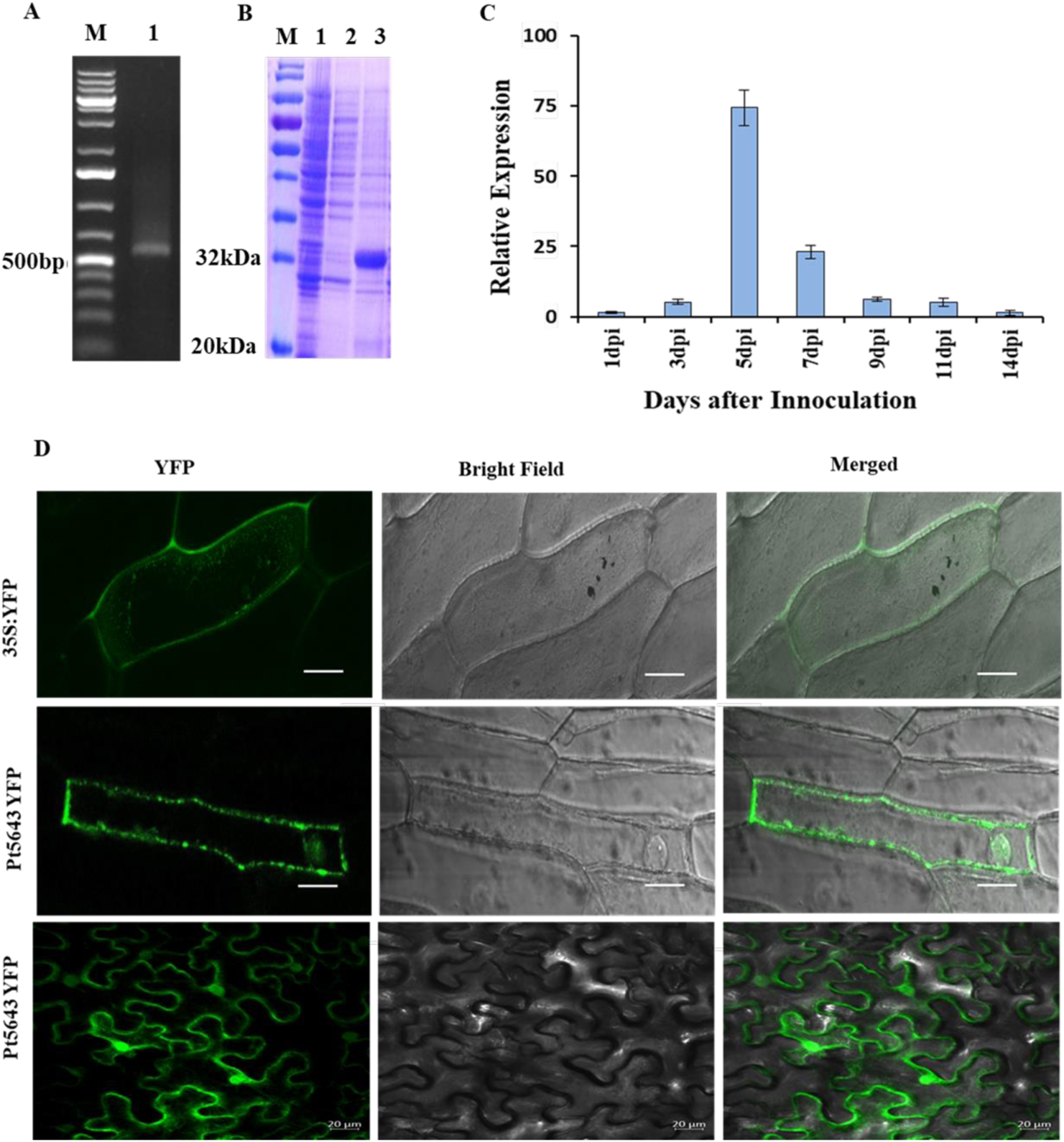
Characterization of *npc2* like gene of *P. triticina* (*Pt5643*) *P. triticina* infected Wheat leaf samples were used for RNA extraction and cDNA synthesis. (A) PCR amplification of Pt5643 gene with a band of 570bp from the infected leaf sample cDNA. (B) Invitro expression analysis of *Pt5643* in 12%SDS-PAGE gel giving a band of 32kDa in pET29a vector. (C) Expression analysis of Pt5643 in the infection cycle. (D) The subcellular localization of Pt5643 in onion epidermal cells (upper two panel) using the particle bombardment method and and *N.benthamiana* (lower panel) using agroinfiltration shows its location in the nucleus and cytoplasm.

The expression of the *Pt5643* gene increased as the infection proceeded up to 5dpi, and after that, started decreasing, suggesting the role of the gene in the course of infection. The subcellular localization of *Pt5643* in onion epidermal cells and *N. benthamiana* suggested it may localize in cytoplasm and nucleus (Figure 5D), suggesting the transport role of *Pt5643* as previously reported by other studies for animals and *Entamoeba histolytica* NPC2 proteins (Bolanos et al., 2016; Frolov et al., 2003; Huang et al., 2007; Zhao et al., 2014).

### 3.9 Functional complementation assay

The complementation assay of the *Pt5643* in the npc2 mutant showed that *Pt5643* could partially, however, significantly complement the yeast *npc2* gene. The *npc2* yeast mutants are reported to show enhanced tolerance against acetic acid treatment, whereas the wild type show reduced growth(Kawahata et al., 2006). The spot assay of *Pt5643* containing npc2 mutants showed reduced growth in the presence of 100mM acetic acids similar to wild-type yeasts. Whereas *npc2* mutants showed significant growth up to(1/1000 dilutions) clearly showing the function complementation of yeast npc2 genes by rust *Pt5643* gene (Fig. 6).

**Fig. 6.**
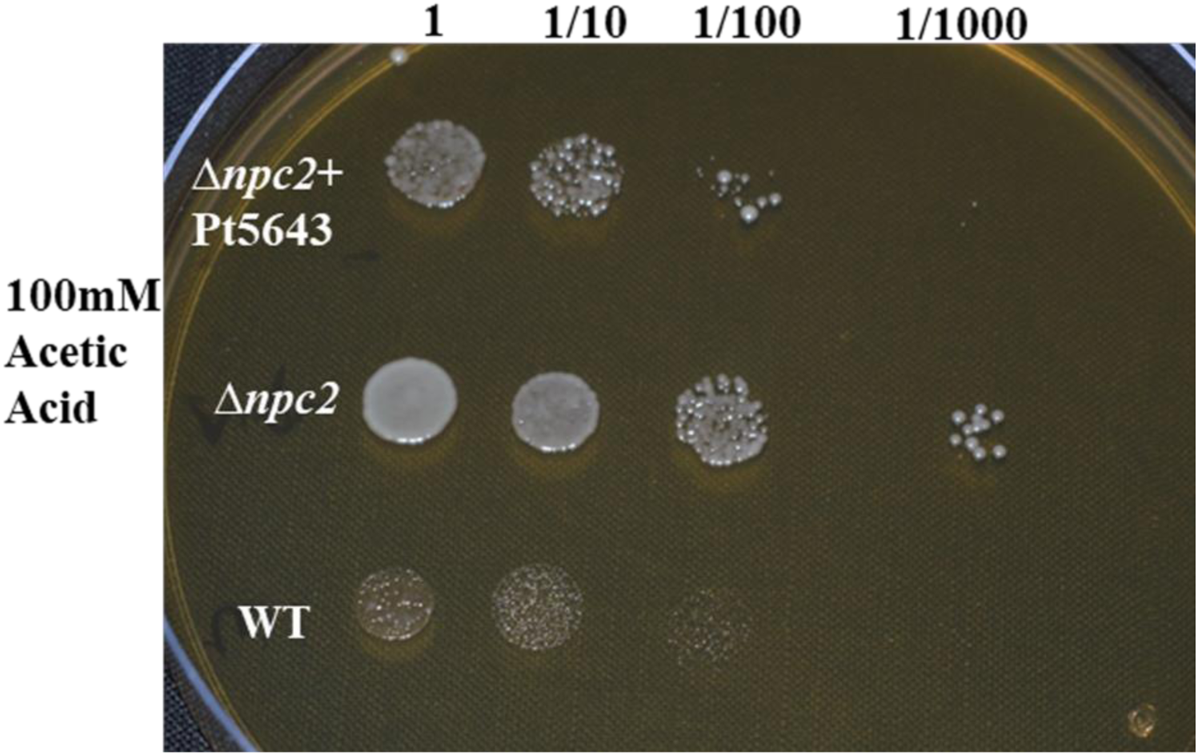
Complementation assay of the *Pt5643* gene in the *npc2* mutant of *S. cerevisiae*. The wild-type yeasts are unable to grow on 100mM acetic acid, while npc2 mutants can grow. The *npc2* mutant containing *Pt5643* shows reduced growths when compared to the npc2 mutant alone, show

### 3.10 Pt5643 transient expression under BAX and H2O2 induced cell death in *N. benthamiana* and *S. cerevisiae*

The transient expression of *Pt5643* in *N. benthamiana* showed *Pt5643* could suppress the cell death induced by BAX gene expression. *Pt5643*, when co-infiltrated with BAX at 0 hrs, failed to hijack the cell death; however, infiltration after BAX after 12hrs did not show any cell death when infiltrated at the same sport where *Pt5643* was infiltrated. Similarly, the infiltration of BAX 24 hrs post infiltration of *Pt5643* also masked cell death (Fig.7) Similar to *N. benthamiana,* the expression of *Pt5643* in yeast revealed its cell death perturbing nature. Diluted yeast cells with 0.0001 OD for wild type*, npc2* mutant yeasts strains, and *Pt5643_npc2* mutants cells when plates on 5mM H2O2 containing plates showed a high number of colonies for the *Pt5643*_*npc2* transformants (approximate 324) than wild type (123) and *npc2* mutant (90) (Fig. 8).

**Fig. 7.**
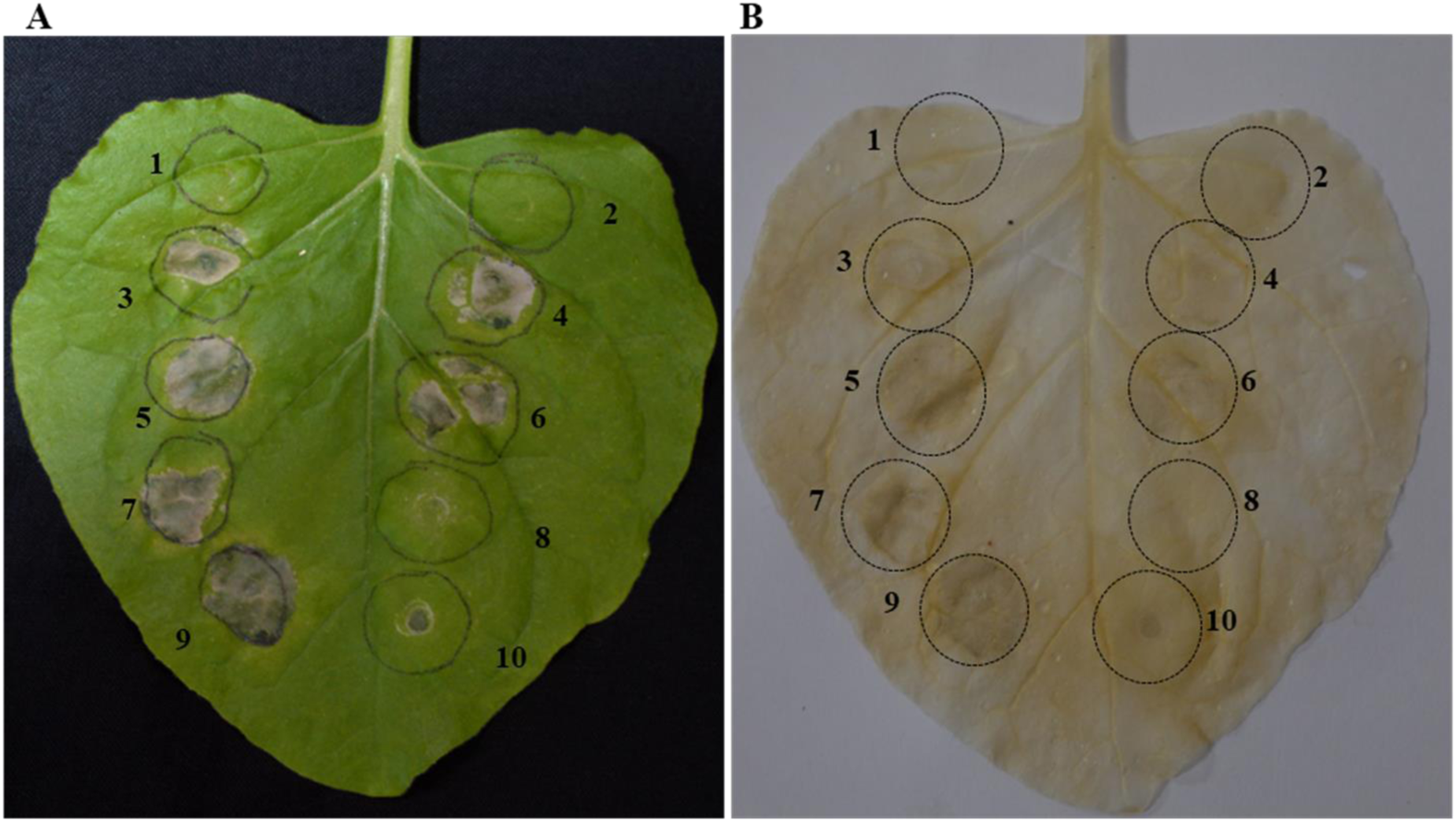
BAX induced cell death suppression assay of *Pt5643* gene in *N. benthamiana* using agroinfiltration method. (A). Infiltration of tobacco leaves and phenotype analysis. (B). Leaf without chlorophyll treated with alcohol and acetic acid mixture (70% ethanol + 10% acetic acid). (**1**.pgwb408-empty vector, **2**.Pt5643_pgwb408, **3**.BAX+ pGWB408, **4**.BAX_pgwb408, **5**.Bax at 0h, **6**.BAX + Pt5643 at 0 h, **7**.Buffer; after 12 h BAX, **8**.Pt5643; after 12 h BAX, **9**. Buffer; after 24 h BAX, **10**. Pt5643; after 24h Bax).

## 4. Discussion

### 4.1 Genome-wide Identification and domain analysis of ML domain superfamily

The *ml/md-2* gene family is present in lower and higher eukaryotes and has been reported to be involved in lipid binding, sterol trafficking, and providing immunity in humans and insects(Liou et al., 2006; Shi et al., 2012; Shimazu et al., 1999). However, the involvement of ML proteins in sterol trafficking and virulence have been reported for few fungal pathogens (Breakspear et al., 2011; Zhao et al., 2014), none of these proteins have been either identified or characterized in plant fungal pathogens. The present investigation pertains to the identification of *ml* gene family members in the 30 different fungal and oomycetes species and further their characterization for host manipulation function. The genome-wide mining identified a total of 84 proteins containing the conserved ML domain, where several proteins harbor other additional domains, including transient receptor potential (TRP) domain, SKP1/BTB/POZ domain, cathepsin propeptide inhibitor domain, and peptidase C1A papain domain. The transient receptor potential (TRP) domain is an integral membrane protein domain present in animals and fungi, where they function as ion channels and are essentially required for the function of intracellular organelles such as vacuoles, endoplasmic reticulum, and lysosomes (Cheng et al., 2010; Puertollano and Kiselyov, 2009). In the present study, 12 out of the 84 identified ML proteins were found to contain the TRP domain and were shown to retain their membrane-bound property (Table 2). In yeast, the TRP protein-encoding gene has been reported to be involved in intracellular calcium ion transport and signaling through the vacuolar membrane (Denis and Cyert, 2002; Palmer et al., 2001). A similar finding has been reported for *Gibberella zeae* (ascomycetes) fungal TRP protein, TRPGz, where TRPY1 knock-out yeast cell expressing fungal TRPGz showing activation under hyperosmolarity, temperature changes, elevated cytosolic Ca^2+^ ion concentration, membrane potential after H_2_O_2_ treatment (Ihara et al., 2013). These previous studies suggest that the TRP domain-containing ML proteins identified in the present study might play a similar role in the environmental stimuli signaling in case plant pathogenic fungi of the class ascomycetes. The presence of cathepsin propeptide inhibitor domain and peptidase C1A papain domain along with ML domain has been found exclusively in oomycetes ML proteins, which suggests functional divergence of these proteins from fungal ML proteins. Intriguingly, in contrast to the investigated fungal species, only members of *Pucciniaceae* have been found to contain NPC2 like domain-containing proteins pointing towards its divergence from the rest of the fungi. Although members of *Clavicipitaceae* have been previously reported to have NCP2 like proteins with lipid-binding function (Zhao et al., 2014), no comparative analysis has been performed across the fungal species.

The identification of ML proteins showed a limited number of ML proteins in fungal species; however, a huge expansion of the ML protein family was observed in mycorrhizal fungus *Rhizophagus irregularis* (Figure 1 B) an obligate biotroph. The fungi and plants are known to contain PG/PI proteins, which are involved in binding with phosphatidylglycerol (PG) and phosphatidylinositol (PI), where PG and PI are important messenger molecules involved in lipid-based signaling (Inohara and Nuñez, 2002). The presence of a high number of *R. irregularis* PG/PI subfamily members suggests that they might be involved in hijacking or interfering with plant lipid-based signaling pathways for successful establishment inside the plant. Previous studies have reported the high expression of the majority of PG/PI genes in *R.irregularis* during infection progression and have also been defined as effectors in *R. clarus*(Chen et al., 2018; Toro and Brachmann, 2016; Zeng et al., 2018). Additionally, multiple secretory nonspecific lipid transfer proteins have been identified, responsible for the transfer of lipids from host to mycorhizal fungi and further recycling (Bago et al., 2002; Jiang et al., 2017; Keymer et al., 2017; Luginbuehl et al., 2017; Zeng et al., 2018). Recent studies have also reported the presence of palmitic acid in the *R. irregularis* hyphae, irrespective of the complete absence of lipid biosynthesis pathway genes, indicating the accumulation of palmitic acid through transport from the host (Wewer et al., 2014).

### 4.2 Phylogenetic analysis of ML domain proteins

The initial studies on the identification of ML gene family in all eukaryotes, including plants, animals, and fungi, identifies only PG/PI type ML proteins in plants and fungi (Inohara and Nuñez, 2002). The absence of NPC2 like proteins in all the majority of fungal species suggests that *npc2* like genes have never been a part of fungal species in due course of evolution. Interestingly, Zhao et al., (Zhao et al., 2014) identified *npc2* like genes in the insect pathogenic fungi (*Metarhizium robertsiii, Metarhizium acridum, and Claviceps pupurea),* encoding NPC2 proteins and shown to exhibit lipid-binding activity similar to aminal NPC2 proteins. Zhao et al., found these genes to be horizontally transferred from the host insect of *Clavicipitaceae*. However, no studies was previously performed since then to identify ML domain (*npc2* like genes) genes in plant fungal pathogens or any other fungal species. The phylogetic analysis of 68 different eukaryotes (Fungi-22, Plants-6, oomycetes-5, protists-28, animals-5) clearly showed that most of fungi contain PG/PI members and clutster separately from animal NPC2, however members from the *Pucciniaceae* (6 species) and previously reported HGT candidate (*C. pupurea*, Cpurp_M1W4K5) from family *Clavicipitaceae* clearly clustered with animal ML proteins especially with insect NPC2 members suggesting their similarity to animal NPC2 than fungal ML proteins (PG/PI).

The similarity of *Pucciniaceae* ML members, with NPC2 like proteins of insects and animals in the phylogenetic analysis, also supported the findings of Zhao et al., 2014 (Fig. 2). Besides, the molecular docking studies also reveal the putative cholesterol binding activity of Puccinia NPC2 proteins, a characteristic of animals and insects NPC2 proteins, thus, indicates their functional conservation across the kingdoms.

An insight into the evolutionary history before diversification revealed that both animals and fungi evolved from the common ancestor “Opisthokonts” and were expected to share the common gene pool. However, the phylogenetic analysis showed the complete absence of the NPC2 class of ML proteins in all the 22 major plant pathogens studied (17 fungal pathogens and 5 oomycetes). From the evolutionary point of view, there are very fewer chances that selection pressure simultaneously led to the loss of genes in multiple organisms. So, it could be possible that identified fungal *npc2* like genes have been evolved independently from plant/fungi-specific *PG/PI* genes in these fungal species. However, the chances of *npc2* evolution in *Puccinia* spp. from *pg/pi* is highly unlikely since none of the *Puccinia* spp. (six spp.) were found to contain *pg/pi* genes (even with the BLAST hit score). Therefore, one of the possible mechanisms of acquisition of *npc2* by *Puccinia* spp. could be horizontal gene transfer (HGT) from animals, insects, or other fungi containing *npc2* genes. Since *Puccinia* spp. and *Claviceps* spp. both infect the wheat plant; therefore, there may be chances of *npc2* like genes getting transferred to *Puccinia* spp. From *Claviceps* spp. However, the *npc2* genes of *Clavicipitaceae* members, including *Claviceps*, clustered separately in the phylogeny away from the *Puccinia* spp., (Figure 2, Figure S2 and S3) suggesting its independent origin, rather than from *Claviceps* spp.

Since the members of *Clavicipitaceae* are entomopathogenic fungi and are known to interact with insects, a previous report for *Clavicipitaceae* members, *C. pupurea,* and *Metarhizium* spp. suggested the origin of *npc2* like genes in these fungi through HGT from host insects (Zhao et al., 2014). Unlike *Clavicipitaceae*, the rust fungal pathogens (*Puccinia* spp.) are not known to feed on the insects; however, there are examples where rust fungi interact with insects in the environment. The interaction of rust fungi and insects may occur through either a vector-host system for spore transmission (Stephanie et al., 2001; Wandeler and Bacher, 2006) or fungal spore infestation. The rust spore infestation by insects of the *Diptera* class is a widespread and frequent phenomenon (Henk et al., 2011), suggesting that *Puccinia* spp. *npc2* genes might also be originated from insects through the HGT event, similar to that of *Clavicipitaceae* members. The hypothesis that *Puccinia* spp. *npc2* genes could be an HGT event from insects appears to be more valid as *Drosophila* NPC2 proteins were clustered together with NPC2 proteins of *Puccinia* spp (Fig. 2). Also, the significant BLASTp score with more than 30% similarity of these clustered NPC2 proteins, and similar 60-64% GC content of encoding genes (45-50% GC content for the rest of the ML gene family members), suggests that members of the *Diptera* family could be the possible donor insect species for *Puccinia npc2* genes.

Moreover, the *npc2* genes have also been identified in few other fungal species of genus *viz Kwoniella spp., P.chlamydosporia,* and *Ustilaginoidea virens,* where all of these fungal species are known to interact either with insects or nematodes in the environment. Therefore, the HGT from insects could be the only possible mechanism to explain the origin of *npc2* genes in these fungal species. However, experimental validation is required to negate neo-functionalization and to provide supporting evidence for HGT events.

### 4.3 3D-Structure prediction, molecular docking, and selection pressure analysis

The molecular docking revealed that three NPC2 like proteins of *Puccinia spp* (Pt5643, P_striiiformis_12, Psorghi_66), and one each from *C.pupurea* (CpM1WA48) and *B. graminis* (BGA0A078N1M3) exhibit binding with cholesterol well as with various sterols like ergosterol, lanosterol, fungisterol, etc (Table 2). Previously it was reported that NPC-2 proteins have the capacity to bind with various sterols because of basic amino acid clusters inside the deep hydrophobic pocket (Liou et al., 2006). The 3D structure analysis of NPC2 like proteins identified in the present study also revealed the presence of basic amino acids (valine, isoleucine, alanine, and phenylalanine) containing deep hydrophobic pockets able to bind with diverse sterols with a high docking score. The superimposition analysis and low Root-Mean-Square Deviation RMSD score (1-2) of these proteins during superimposition with bovine NPC-2 protein crystal structure (data not shown) suggest a functional similarity of these proteins with animal NPC-2 proteins. Besides, all the five proteins also showed significant docking scores with isopentenyl pyrophosphate (IPP), geranyl pyrophosphate (GPP), and farnesyl pyrophosphate (FPP) (Table 3), a precursor of various phytoalexins produced against infection of fungi and various pathogens by the plants (Chávez-Moctezuma and Lozoya-Gloria, 1996; Walters et al., 2008).

All the three fungal pathogens (*Puccinia spp*, *C.purpurea,* and *B.graminis*) are biotrophic and dependent on the host for its survival and propagation, so it could be expected that fungal NPC-2 like proteins have retained the conserved functions of lipid trafficking for fungal survival or engaging the sterols or lipids responsible for signaling that could otherwise reveal the presence of pathogen inside the host cell. Therefore, it could be suggested that NPC2 like proteins are involved in processes against the plant defense mechanism. In addition, the value of Ka/Ks for a few of the fungal *npc-2* like genes was found to be less than one. which suggests purifying selection pressure and further corroborate the functional conservation of NPC2 like genes in fungus.

### 4.4 Functional complementation assay

In the case of *npc2* genes, the studies have reported that wild-type yeast show reduced growth in acetic acid, while mutant npc2 strains are resistant to the treatment(Kawahata et al., 2006). The functional complementation of one of the *P.triticina* gene *Pt5643* with yeast *npc2 (*ML homolog) homolog in the presence of acetic acid (100mM) shows that *Pt5643* partially however significantly complements the function npc2 mutant showing phenotype similar to wild type yeast, further suggesting the *Pt5643* to play a role similar to yeasts NPC2 (sterol trafficking). Though the *Pt5643* gene did not complement the function of later completely, it may play a relatively diverged function in *Puccinia* spp.

### 4.5 *Pt5643* transient expression under BAX and H2O2 induced cell death in N. benthamiana and S. cerevisiae

The BAX and H_2_O_2_ induced program cell death (PCD) hijacking capacity of *Pt5643* gene after expressing in *N. benthamiana* and *S. cerevisiae* further suggested its putative function in *P.triticina* as a potential effector. In the case of biotrophic pathogens, hijacking of the host induced cell death by interfering with host immune signaling is one of the effector’s primary functions. Moreover, cholesterol is also known to play a crucial role in BAX-mediated program cell death (PCD)(Lucken-Ardjomande et al., 2008). The cholesterol accumulation in the cell regulates PCD. At low concentrations, cholesterol promotes the BAX-induced cell death, while high concentration inhibits by changing the membrane property(Li et al., 2020; Mignard et al., 2014). The molecular docking of *Pt5643* has shown it to bind cholesterol, and further subcellular localization also showed them to be cytoplasm localized. Therefore, the cell death hijacking function of the *Pt5643* could be due to its cholesterol trafficking and further accumulation in the cell.

The overall results in the study showed that NPC2 like proteins in fungal pathogens might bind to various sterols present in the sterol synthesis pathway, perform trafficking functions, and are probably originated from animal/insect species. Insects are in continuous interaction with fungal pathogens since spores of biotrophic fungal pathogens could be an easy source of nutrition for mycophagous insects. Therefore, the presence of insect-related genes in the *Puccinia* sp, *Claviceps* sp, and *Tremellales* sp might result from the interaction between ancestors of insects and these pathogens and for possible HGT events. Moreover, its presence in a large number of Pucciniaceae members suggests its important role as a conserved effector that hijacks the basic biological process in PCD.

## 5. Conclusions

This study sheds light on the status of the *ml* gene family in plant pathogenic fungi, and its possible role in plant-pathogen interactions. The ML domain is also associated with various other domains and putatively involved in diverse functions. The results also showed that NPC2 like proteins of *Puccinia*ceae, *Claviceptaecace,* and *Tremellaceae* have possible insect origin and are transferred to these pathogens through the HGT mechanism. Most of the *ml* gene family members in fungi are going through purifying selection except a few *B.graminis*, *P.striiformis*, *and R.irregularis ml* genes with strong positive selection pressure. The *npc2*-like genes are already reported to perform sterol transport in humans and other animals, including insects, in association with the *npc1* gene across the membrane. The acquisition of *npc2* like effector genes and expansion in biotrophic pathogens like rust, Claviceps, *R. irregularis* could be associated with their obligate biotrophic nature, as these pathogens hijack the host signaling system to remain undetected and require raw materials to sustain inside the host.

## Acknowledgments

TRS is thankful to the Department of Science and Technology, Govt. of India, for JC Bose National Fellowship. RJ is thankful^i^ to the University Grants Commission (UGC), New Delhi for providing Junior Research Fellowship (JRF).

## CRediT authorship contribution statement

**Rajdeep Jaswal:** Conceptualization, Methodology, Software, Analysis, Data curation, Experiments, Writing - original draft. **Himanshu Dubey:** Formal analysis. **Kanti Kiran**: Formal analysis. **Hukam Rawal:** Formal analysis. **Sivasubramanian Rajarammohan:** Formal analysis. **Humira Sonah:** Formal analysis. **Rupesh Deshmukh**- Formal analysis. **Pramod Prasad**: resource**. Subhash C Bhardwaj**: resource. **Naveen Gupta-** Supervision. **Tilak Raj Sharma:** Conceptualization, Methodology, Supervision, Funding acquisition, writing-review, and editing.

## References

Altschul, S.F., Gish, W., Miller, W., Myers, E.W., Lipman, D.J., 1990. Basic local alignment search tool. J. Mol. Biol. 215, 403–410.

Ao, J., Ling, E., Rao, X., Yu, X.-Q., 2008. A novel ML protein from *Manduca sexta* may function as a key accessory protein for lipopolysaccharide signaling. Mol. Immunol. 45, 2772–2781.

Bago, B., Zipfel, W., Williams, R.M., Jun, J., Arreola, R., Lammers, P.J., Pfeffer, P.E., Shachar-Hill, Y., 2002. Translocation and utilization of fungal storage lipid in the arbuscular mycorrhizal symbiosis. Plant Physiol. 128, 108–124.

Bikadi, Z., Hazai, E., 2009. Application of the PM6 semi-empirical method to modeling proteins enhances docking accuracy of AutoDock. J. Cheminform. 1, 15.

Bolanos, J., Betanzos, A., Javier-Reyna, R., Garcia-Rivera, G., Huerta, M., Pais-Morales, J., Gonzalez-Robles, A., Rodriguez, M.A., Schnoor, M., Orozco, E., 2016. EhNPC1 and EhNPC2 proteins participate in trafficking of exogenous cholesterol in *Entamoeba histolytica* trophozoites: relevance for phagocytosis. PLoS Pathog. 12, e1006089.

Breakspear, A., Pasquali, M., Broz, K., Dong, Y., Kistler, H.C., 2011. Npc1 is involved in sterol trafficking in the filamentous fungus *Fusarium graminearum*. Fungal Genet. Biol. 48, 725–730.

Chávez-Moctezuma, M.P., Lozoya-Gloria, E., 1996. Biosynthesis of the sesquiterpenic phytoalexin capsidiol in elicited root cultures of chili pepper (*Capsicum annuum*). Plant Cell Rep. 15, 360–366.

Chen, E.C.H., Morin, E., Beaudet, D., Noel, J., Yildirir, G., Ndikumana, S., Charron, P., St-Onge, C., Giorgi, J., Krüger, M., 2018. High intraspecific genome diversity in the model arbuscular mycorrhizal symbiont *Rhizophagus irregularis*. New Phytol. 220, 1161–1171.

Cheng, X., Shen, D., Samie, M., Xu, H., 2010. Mucolipins: Intracellular TRPML1-3 channels. FEBS Lett. 584, 2013–2021.

Coelho, M.A., Gonçalves, C., Sampaio, J.P., Gonçalves, P., 2013. Extensive intra-kingdom horizontal gene transfer converging on a fungal fructose transporter gene. PLoS Genet. 9, e1003587.

Denis, V., Cyert, M.S., 2002. Internal Ca2+ release in yeast is triggered by hypertonic shock and mediated by a TRP channel homologue. J Cell Biol 156, 29–34.

Dong, Y., Aguilar, R., Xi, Z., Warr, E., Mongin, E., Dimopoulos, G., 2006. Anopheles gambiae immune responses to human and rodent Plasmodium parasite species. PLoS Pathog 2, e52.

Fitzpatrick, D.A., 2012. Horizontal gene transfer in fungi. FEMS Microbiol. Lett. 329, 1–8.

Frolov, A., Zielinski, S.E., Crowley, J.R., Dudley-Rucker, N., Schaffer, J.E., Ory, D.S., 2003. NPC1 and NPC2 regulate cellular cholesterol homeostasis through generation of LDL-derived oxysterols. J. Biol. Chem.

Gruber, A., Manček, M., Wagner, H., Kirschning, C.J., Jerala, R., 2004. Structural model of MD-2 and functional role of its basic amino acid clusters involved in cellular lipopolysaccharide recognition. J. Biol. Chem. 279, 28475–28482.

Halgren, T.A., 1996. Merck molecular force field. I. Basis, form, scope, parameterization, and performance of MMFF94. J. Comput. Chem. 17, 490–519.

Henk, D.A., Farr, D.F., Aime, M.C., 2011. Mycodiplosis (Diptera) infestation of rust fungi is frequent, wide spread and possibly host specific. Fungal Ecol. 4, 284–289.

Huang, X., Warren, J.T., Buchanan, J., Gilbert, L.I., Scott, M.P., 2007. Drosophila Niemann-Pick type C-2 genes control sterol homeostasis and steroid biosynthesis: a model of human neurodegenerative disease. Development 134, 3733–3742.

Ihara, M., Hamamoto, S., Miyanoiri, Y., Takeda, M., Kainosho, M., Yabe, I., Uozumi, N., Yamashita, A., 2013. Molecular bases of multimodal regulation of a fungal transient receptor potential (TRP) channel. J. Biol. Chem. 288, 15303–15317.

Inohara, N., Nuñez, G., 2002. ML–a conserved domain involved in innate immunity and lipid metabolism. Trends Biochem. Sci. 27, 219–221.

Jaswal, R., Dubey, H., Kiran, K., Singh, P.K., Rawal, H.C., Bhardwaj, S.C., Prasad, P., Gupta, N., Sharma, T.R., 2019. Comparative secretome analysis of Indian wheat leaf rust pathogen *Puccinia triticina*. Indian J. Agric. Sci. 89, 1688–1692.

Jaswal, R., Kiran, K., Rajarammohan, S., Dubey, H., Singh, P.K., Sharma, Y., Deshmukh, R., Sonah, H., Gupta, N., Sharma, T.R., 2020a. Effector Biology of Biotrophic Plant Fungal Pathogens: Current Advances and Future Prospective. Microbiol. Res. 126567.

Jaswal, R., Rajarammohan, S., Dubey, H., Sharma, T.R., 2020b. Smut fungi as a stratagem to characterize rust effectors: opportunities and challenges. World J. Microbiol. Biotechnol. 36, 1–10.

Jiang, Y., Wang, W., Xie, Q., Liu, N., Liu, L., Wang, D., Zhang, X., Yang, C., Chen, X., Tang, D., 2017. Plants transfer lipids to sustain colonization by mutualistic mycorrhizal and parasitic fungi. Science (80-.). eaam9970.

Johannessen, B.R., Skov, L.K., Kastrup, J.S., Kristensen, O., Bolwig, C., Larsen, J.N., Spangfort, M., Lund, K., Gajhede, M., 2005. Structure of the house dust mite allergen Der f 2: Implications for function and molecular basis of IgE cross-reactivity. FEBS Lett. 579, 1208–1212.

Jupatanakul, N., Sim, S., Dimopoulos, G., 2014. *Aedes aegypti* ML and Niemann-Pick type C family members are agonists of dengue virus infection. Dev. Comp. Immunol. 43, 1–9.

Juretic, N., Hoen, D.R., Huynh, M.L., Harrison, P.M., Bureau, T.E., 2005. The evolutionary fate of MULE-mediated duplications of host gene fragments in rice. Genome Res. 15, 1292–1297.

Kawahata, M., Masaki, K., Fujii, T., Iefuji, H., 2006. Yeast genes involved in response to lactic acid and acetic acid: acidic conditions caused by the organic acids in *Saccharomyces cerevisiae* cultures induce expression of intracellular metal metabolism genes regulated by Aft1p. FEMS Yeast Res. 6, 924–936.

Keymer, A., Pimprikar, P., Wewer, V., Huber, C., Brands, M., Bucerius, S.L., Delaux, P.-M., Klingl, V., von Roepenack-Lahaye, E., Wang, T.L., 2017. Lipid transfer from plants to arbuscular mycorrhiza fungi. Elife 6, e29107.

Kim, H.M., Park, B.S., Kim, J.-I., Kim, S.E., Lee, J., Oh, S.C., Enkhbayar, P., Matsushima, N., Lee, H., Yoo, O.J., 2007. Crystal structure of the TLR4-MD-2 complex with bound endotoxin antagonist Eritoran. Cell 130, 906–917.

Kiran, K., Rawal, H.C., Dubey, H., Jaswal, R., Bhardwaj, S.C., Prasad, P., Pal, D., Devanna, B.N., Sharma, T.R., 2017. Dissection of genomic features and variations of three pathotypes of *Puccinia striiformis* through whole genome sequencing. Sci. Rep. 7.

Kiran, K., Rawal, H.C., Dubey, H., Jaswal, R., Devanna, B.N., Gupta, D.K., Bhardwaj, S.C., Prasad, P., Pal, D., Chhuneja, P., 2016. Draft genome of the wheat rust pathogen (*Puccinia triticina*) unravels genome-wide structural variations during evolution. Genome Biol. Evol. 8, 2702–2721.

Kumar, S., Stecher, G., Tamura, K., 2016. MEGA7: Molecular Evolutionary Genetics Analysis version 7.0 for bigger datasets. Mol. Biol. Evol. msw054.

Li, K., Deng, Y., Deng, G., Chen, P., Wang, Y., Wu, H., Ji, Z., Yao, Z., Zhang, X., Yu, B., 2020. High cholesterol induces apoptosis and autophagy through the ROS-activated AKT/FOXO1 pathway in tendon-derived stem cells. Stem Cell Res. Ther. 11, 1–16.

Li, W.-H., Gojobori, T., Nei, M., 1981. Pseudogenes as a paradigm of neutral evolution. Nature 292, 237–239.

Liao, J.-X., Yin, Z.-X., Huang, X.-D., Weng, S.-P., Yu, X.-Q., He, J.-G., 2011. Cloning and characterization of a shrimp ML superfamily protein. Fish Shellfish Immunol. 30, 713–719.

Librado, P., Rozas, J., 2009. DnaSP v5: a software for comprehensive analysis of DNA polymorphism data. Bioinformatics 25, 1451–1452.

Liou, H.-L., Dixit, S.S., Xu, S., Tint, G.S., Stock, A.M., Lobel, P., 2006. NPC2, the protein deficient in Niemann-Pick C2 disease, consists of multiple glycoforms that bind a variety of sterols. J. Biol. Chem. 281, 36710–36723.

Livak, K.J., Schmittgen, T.D., 2001. Analysis of relative gene expression data using real-time quantitative PCR and the 2− ΔΔCT method. methods 25, 402–408.

Lucken-Ardjomande, S., Montessuit, S., Martinou, J.-C., 2008. Bax activation and stress-induced apoptosis delayed by the accumulation of cholesterol in mitochondrial membranes. Cell Death Differ. 15, 484–493.

Luginbuehl, L.H., Menard, G.N., Kurup, S., Van Erp, H., Radhakrishnan, G. V, Breakspear, A., Oldroyd, G.E.D., Eastmond, P.J., 2017. Fatty acids in arbuscular mycorrhizal fungi are synthesized by the host plant. Science (80-.). eaan0081.

Mignard, V., Lalier, L., Paris, F., Vallette, F.M., 2014. Bioactive lipids and the control of Bax pro-apoptotic activity. Cell Death Dis. 5, e1266–e1266.

Palmer, C.P., Zhou, X.-L., Lin, J., Loukin, S.H., Kung, C., Saimi, Y., 2001. A TRP homolog in Saccharomyces cerevisiae forms an intracellular Ca2+-permeable channel in the yeast vacuolar membrane. Proc. Natl. Acad. Sci. 98, 7801–7805.

Puertollano, R., Kiselyov, K., 2009. TRPMLs: in sickness and in health. Am. J. Physiol. Physiol. 296, F1245–F1254.

Qiu, H., Cai, G., Luo, J., Bhattacharya, D., Zhang, N., 2016. Extensive horizontal gene transfers between plant pathogenic fungi. BMC Biol. 14, 41.

Shi, X.-Z., Zhong, X., Yu, X.-Q., 2012. Drosophila melanogaster NPC2 proteins bind bacterial cell wall components and may function in immune signal pathways. Insect Biochem. Mol. Biol. 42, 545–556.

Shimazu, R., Akashi, S., Ogata, H., Nagai, Y., Fukudome, K., Miyake, K., Kimoto, M., 1999. MD-2, a molecule that confers lipopolysaccharide responsiveness on Toll-like receptor 4. J. Exp. Med. 189, 1777–1782.

Slot, J.C., Rokas, A., 2011. Horizontal transfer of a large and highly toxic secondary metabolic gene cluster between fungi. Curr. Biol. 21, 134–139.

Stephanie, K., Andreas, K., Teja, T., 2001. Interactions between the rust fungus *Puccinia punctiformis* and ectophagous and endophagous insects on creeping thistle. J. Appl. Ecol. 38, 548–556.

Toro, K.S., Brachmann, A., 2016. The effector candidate repertoire of the arbuscular mycorrhizal fungus *Rhizophagus clarus*. BMC Genomics 17, 101.

Trifinopoulos, J., Nguyen, L.-T., von Haeseler, A., Minh, B.Q., 2016. W-IQ-TREE: a fast online phylogenetic tool for maximum likelihood analysis. Nucleic Acids Res. 44, W232–W235.

Vizzini, A., Bonura, A., Longo, V., Sanfratello, M.A., Parrinello, D., Cammarata, M., Colombo, P., 2015. Isolation of a novel LPS-induced component of the ML superfamily in Ciona intestinalis. Dev. Comp. Immunol. 53, 70–78.

Walters, D., Newton, A.C., Lyon, G., 2008. Induced resistance for plant defence: a sustainable approach to crop protection. John Wiley & Sons.

Wandeler, H., Bacher, S., 2006. Insect-transmitted urediniospores of the rust *Puccinia punctiformis* cause systemic infections in established Cirsium arvense plants. Phytopathology 96, 813–818.

Wewer, V., Brands, M., Dörmann, P., 2014. Fatty acid synthesis and lipid metabolism in the obligate biotrophic fungus *Rhizophagus irregularis* during mycorrhization of L otus japonicus. Plant J. 79, 398–412.

Xu, D., Zhang, Y., 2011. Improving the physical realism and structural accuracy of protein models by a two-step atomic-level energy minimization. Biophys. J. 101, 2525–2534.

Xu, S., Benoff, B., Liou, H.-L., Lobel, P., Stock, A.M., 2007. Structural basis of sterol binding by NPC2, a lysosomal protein deficient in Niemann-Pick type C2 disease. J. Biol. Chem. 282, 23525–23531.

Yang, J., Yan, R., Roy, A., Xu, D., Poisson, J., Zhang, Y., 2015. The I-TASSER Suite: protein structure and function prediction. Nat. Methods 12, 7–8.

Yin, Z., Zhu, B., Feng, H., Huang, L., 2016. Horizontal gene transfer drives adaptive colonization of apple trees by the fungal pathogen Valsa mali. Sci. Rep. 6, 33129.

Zeng, T., Holmer, R., Hontelez, J., te Lintel-Hekkert, B., Marufu, L., de Zeeuw, T., Wu, F., Schijlen, E., Bisseling, T., Limpens, E. 2018. Host-and stage-dependent secretome of the arbuscular mycorrhizal fungus *Rhizophagus irregularis*. Plant J. 94, 411–425.

Zhao, H., Xu, C., Lu, H.-L., Chen, X., Leger, R.J.S., Fang, W., 2014. Host-to-pathogen gene transfer facilitated infection of insects by a pathogenic fungus. PLoS Pathog 10, e1004009.

